# Exploring the utility of regulatory network-based machine learning for gene expression prediction in maize

**DOI:** 10.1101/2023.05.11.540406

**Authors:** Taylor H. Ferebee, Edward S. Buckler

## Abstract

Genomic selection and gene editing in crops could be enhanced by multi-species, mechanistic models predicting effects of changes in gene regulation. Current expression abundance prediction models require extensive computational resources, hard-to-measure species-specific training data, and often fail to incorporate data from multiple species. We hypothesize that gene expression prediction models that harness the regulatory network structure of *Arabidopsis thaliana* transcription factor-target gene interactions will improve on the present maize models. To this end, we collect 147 *Oryza sativa* and 99 *Sorghum bicolor* gene expression assays and assign them to maize family-based orthologous groups. Using three popular graph-based machine learning frameworks, including a shallow graph convolutional autoencoder, a deep graph convolutional autoencoder, and the inductive GraphSage strategy, we encode an *Arabidopsis thaliana* integrated gene regulatory network (iGRN) structure and TF gene expression values to predict gene expression both within and between species. We then evaluate the network methods against a partial least-squares baseline. We find that the baseline gives the best predictions within species, with Spearman correlations averaging between 0.74 and 0.78. The graph autoencoder methods were more variable with correlations between -0.1 and 0.65. In particular, the GraphSage and deep autoencoders performed the worst, and the shallow autoencoders performed the best. In the most challenging prediction context, where predictions were in new species and on genes that were not seen, we found that the shallow graph autoencoder framework averaged around 0.65. Unlike initial thoughts about preserved network structure improving gene expression predictions, this study shows that within-species predictions only need simple models, such as partial least squares, to capture expression variations. In cross-species predictions, the best model is often a more complex strategy utilizing regulatory network structure and other studies’ expressions.

## Introduction

In recent years, gene expression prediction models have been introduced as a computational tool to identify key regulatory elements that influence the effects of disease variants [1] or describe the nature of regulatory grammars [2–4]. Because of the ability of the plant genetics community to study variations of complex traits, understand the evolutionary processes behind these traits, and target sites for gene editing, expression prediction holds a lot of research potential. To gather the necessary resolution to explore these concepts, researchers generally focus on the systematic role of regulatory machinery that facilitates variation in gene expression over space and time [5, 6]. Gene regulation requires elaborate interplay between linked (*cis*) and distant (*trans*) regulatory elements. In the *cis*-case, DNA sequence-specific transcription factors bind to the DNA to modulate nearby gene expression In addition to transcription factors, untranslated regions of RNA modulate gene expression through length variation and through offering variants of mRNA from the same gene [7]. In plants, mutations in these *cis*-regulatory regions contribute to domestication [8, 9], adaptation [10, 11], and, in some cases, are the target of allele-generative pursuits in next-generation breeding approaches [12, 13].

Across plant species, the relationship between genes and transcription factors has been one of the main focus for understanding the transcriptomic patterning. For example, just under 50% of orthologous genes between *Arabidopsis thaliana* and *Oryza sativa*, species that diverged about 150 million years ago [14], show similar transcriptomic responses while under abiotic stress [15, 16]. Further, the different responses were due to changes in regulatory elements, such as transcription factor expression. It has also been shown that tissue-specific transcription factor activity induces patterning in gene target specificity [17]. Due to the multidimensional relationship between tissue, experimental conditions, and species, few studies have attempted to model gene expression patterns between species.

Gene regulatory networks (GRNs) represent the emergence of transcriptomic patterning by providing a graph representation of the TF to target gene relationships. GRNs have been crucial to understanding not only the interactions between genes, but also the coordinated targeting of specific genes by multiple transcription factors [18, 19]. Assays, such as CHIP-seq and ATAC-seq, inform the majority of the transcription factor-gene relationships in these networks, and with sufficient experimental data, these networks can also integrate the 3-D chromatin structure and dynamics to reveal complex, time-sensitive, and species-specific players in gene regulation [20–23]. Unfortunately, these combinations of experiments can be expensive, hard to measure, and computationally difficult to utilize.

There have been several recent successes in overcoming the limitations of predicting gene expressions or other large-scale datasets using deep learning. Generally, deep learning refers to computational methods that aim to learn a hierarchical representation of data by functionally relating the data in many layers [24]. In plant data, these models have learned to predict chromatin state from sequence [25], identify and classify key stress-responsive genes [26], and detect seasonal changes across fields [27]. In addition to these successes, many state-of-the-art models take in not only numerical data but also the structure of the relationships between the data. Traditionally, non-Euclidean graphical structures represent these complex, multidimensional relationships. The models, graph neural networks, use the same general machinery that many deep learning methods use, but take special care to ensure that the representation of the data is preserved through each abstraction. Graph neural networks have shown particular success in exploring disease and pest relationships in *Arabidopsis thaliana*, inferring protein-protein interactions key to biological functions [28], and finally, inferring gene expression in mice, given co-expression and gene interaction networks [29].

Graph neural networks solve problems typically associated with complex, non-Euclidean dimension reduction, image extraction, and link or node prediction. Strategies to solve these problems can be categorized into spectral and spatial-based approaches. Spectral approaches, such as the common graph convolutional network (GCN), focus primarily on the graph spectrum, or the set of eigenvectors and eigenvalues, associated with the adjacency matrix of the graph [30]. In these cases, the graph spectrum is used to map its structure to a Euclidean space for which convolutional operations can be performed similarly to convolutional neural networks. Spatial approaches work at the node and neighboring node-edge level to aggregate information across nodes. Due to this aggregation, many special methods are amenable to large networks. For example, GraphSage [31], aggregates node data in order to create low dimensional representations of unseen data. This aggregation to embedding step makes GraphSage a natural choice for large biological studies, and researchers use the method to explore large-scale predictions for large-scale protein-protein interactions [32] and drug-disease relationships [33].

However, to create the models, care must be taken to preserve the broad evolutionary relationships. Plant systems, because of genome duplication and adaptive responses to selection pressures, generally have larger expansion of transcription factor families than animal ones [34]. These expansions, however, often act in parallel, which preserves researchers’ aim of accurate and precise global quantification of TF-gene relationships. In *Arabidopsis thaliana*, researchers mined thousands of experimental and computational data to find the most likely interactions between genes and transcription factors [35]. Their goal was to use machine learning machinery to infer a network that was biologically supported and could be used to functionally validate biological hypotheses. The resulting network identified key regulators of reactive oxygen stress response [35]. In a cross-species comparison of gene regulatory networks, researchers inferred stress-responsive sub-networks that are evolutionarily conserved between *Marchantia polymorpha* and *Arabidopsis thaliana*, allowing for new hypotheses and methods that can provide insight into divergent species’ gene regulatory mechanisms [36]. In this work, we use popular network embedding-based deep learning methods to predict gene expression in maize experiments. We design the machine learning task around the embedding of a regulatory network structure from the model species, *Arabidopsis thaliana*. Upon training the model on *Zea mays, Arabidopsis thaliana*, and *Sorghum bicolor*, we observe improved correlations between predictions and observed values relative to previous regression and machine learning models. Finally, we examine the characteristics of the regulatory network that may affect the prediction of gene expression.

## Materials and methods

### Raw data and preprocessing

Gene expression data for *Zea mays, Sorghum bicolor*, and *Oryza sativa* was downloaded and processed as in [37]. In summary, datasets were downloaded from the National Center for Biotechnology Information (NCBI) Sequence Read Archive (SRA), trimmed and quality checked with FastQC and Sickle, and aligned to Maize V4, Sorghum 3.1.1, and *Oryza sativa* V7 genomes with HISAT2 [37], and read counts were normalized to transcripts per million with Stringtie. Identifications of transcription factors for *Zea mays, Sorghum bicolor*, and *Oryza sativa* were downloaded from the Plant Transcription Factor Database website [38]. Orthologous groups based on gene families were recovered from the PLAZA 4.5 database [39]. The Arabidopsis integrated gene regulatory network (iGRN) was downloaded from the supplementary of [35]. The handling of the data sets was done with custom R scripts that utilize the data.table and dplyr packages [40] were used to process the data sets. All data, including code to generate input formats, are enclosed in the code repository on Zenodo (doi: 10.5281/zenodo.7194372). Supplemental Figure 1 shows the flow chart of the data preprocessing.

### Orthogroup assignments

For each species, gene expression matrices were transformed into orthogroup matrices by cross-referencing the PLAZA gene family-based orthogroups [39]. To create an orthogroup expression matrix, the mean orthogroup expression per experiment was taken across replicates. Expression levels were log-transformed and normalized by the standard deviation of an orthogroup’s expression across experiments. The Arabidopsis iGRN nodes were similarly transformed into orthogroups. The nodes and their corresponding edges were collapsed to create a network with orthogroups as the nodes rather than the original Arabidopsis genes. The resulting network contained 756 transcription factors and 18771 target genes. From this list, an adjacency matrix was created to represent the network’s connections.

### Model splitting

Training and testing splits were made at the orthogroup and experimental levels. In the case we predict expression on orthogroups that the model has seen, we perform an 80:20 train-test split across experiments. To ensure there is minimal experiment-experiment leakage, we completely mask these values during the training process. In the case we predict expression on genes that have not been seen by the model, we do a two part procedure for training. We first perform an 80:20 split on the experiments. Then, we perform another 80:20 split on the orthogroups. The final predictions are completed on the unseen genes within the test experiments.

### Model development

Expression abundance predictions were made with two architectures, structural embedding with random forest (graph autoencoder models) [33] and a partial least squares model. All models structural embedding models were constructed using Python 3 with the Pytorch Geometric [41] and Scikit-learn libraries. Models were tested in 4 scenarios: 1) within-species prediction of the unseen expression profile when all genes have been seen by the model, 2) prediction in maize from a single species when all genes have been seen by the model, 3) prediction of unseen expression profile when all genes have not been seen by the model, and 4) prediction of maize expression profiles from all species.

Expression predictions were done with two architectures: structural embedding with random forest prediction layer (graph autoencoder models) and a baseline partial least squares model. We tested three main architectures for the GAE-a shallow graph convolutions, deep graph convolutions, and Graph Sage. The random forest layer was constructed with 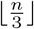 decision trees, as suggested by [42]. All structural embedding models were constructed in Python 3 with the Pytorch Geometric library [41]. Predictions were made within each related species and into maize (target species). We tested both the PLS and structural embedding random forest architectures on a new test set containing genes which the model has been trained on. In structural embedding models, we also tested in situations where the predicted genes’ expression profiles are masked during training (Supplementary Figure 1).

### Partial least squares baseline

To establish a baseline performance, we use a partial least squares model. In this case, consider **X**_*nonT F*_ to be a gene expression matrix consisting of orthogroups not associated with transcription factors. Similarly, let **X**_*T F*_ be a gene expression matrix of only transcription factor orthogroups. We want to predict each column of the non-transcription factor, **X**_*nonT F*_, from a combination of the columns of **X**_*T F*_. Partial least squares transforms the space of predictors in **X**_*T F*_ to orthogonal variables, which we then use for prediction (Fig1). We chose to use the PLS2 algorithm which both **Y**’s and **X**_*T F*_ s variable spaces are simultaneously modeled. Here, the underlying model, with components *T* and *U*, loading *P* and *Q*, and error terms *E* and *F* becomes,

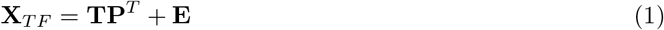

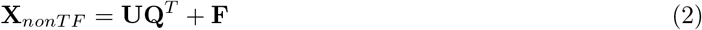

**Fig 1.**
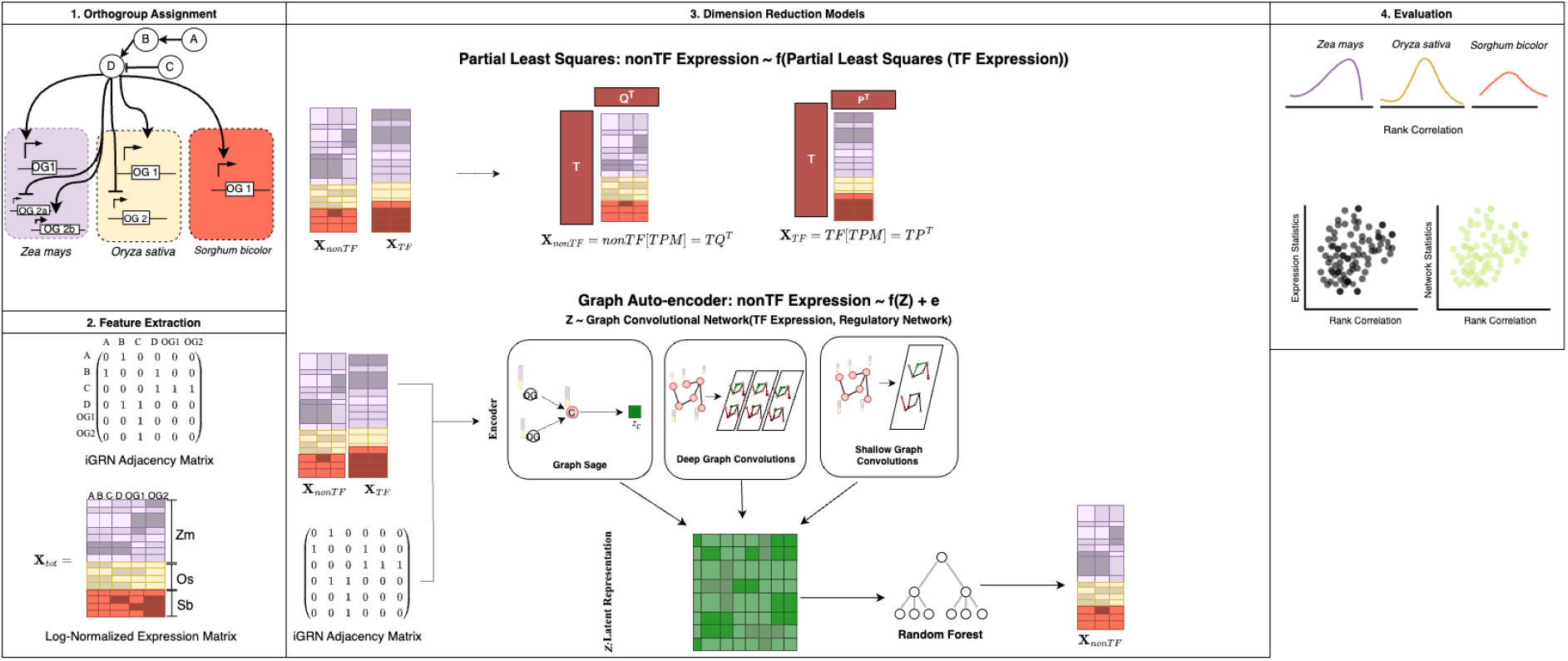
Schematic of Cross Species Prediction Model. Flowchart of predicting cross-species gene expression in maize. First, we assign orthogroups to each of the species, *Zea mays*, and *Sorghum bicolor*. Then, we gather species’ gene expression data and the Arabidopsis integrated gene regulatory network(iGRN). The network is converted to an adjacency matrix, and species expression data is log-normalized and transformed into an orthogroup-by-condition expression matrix. Next we establish a baseline prediction model using partial least squares. We then construct predictions from three different encoding strategies for the encoding of the expression and network– Shallow Graph Convolutions, Deep Graph Convolutions, and Graph Sage. The resulting latent representation is then used in a random forest model in order to predict unseen expressions across various conditions. Finally, we evaluate the models by examining the Spearman correlation between observed and predicted expression profiles. To determine the drivers of predictions, we also examine expression statistics versus model performance and network statistics versus model performance relationships.

**Fig 2.**
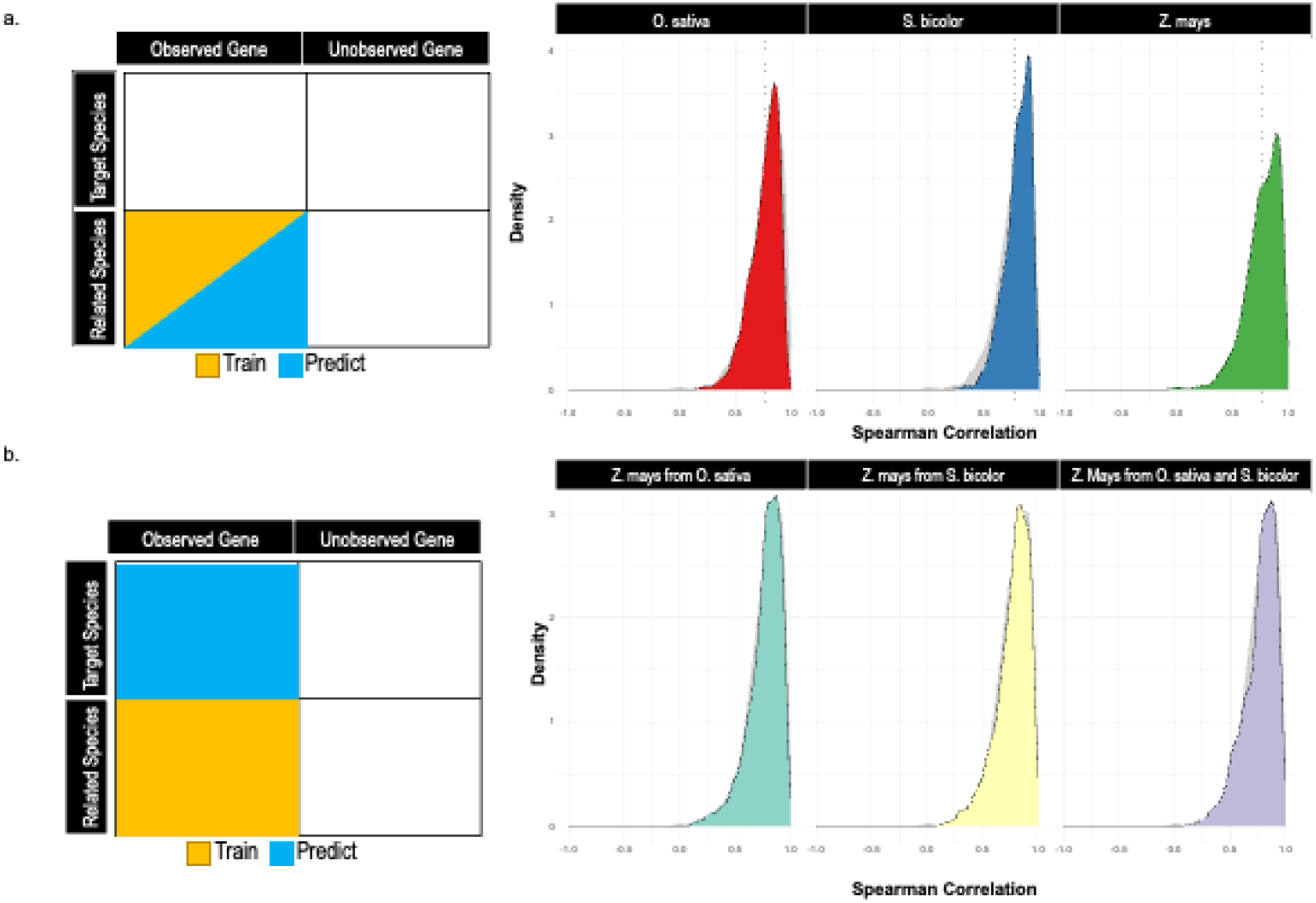
Baseline partial least squares prediction performance of within-species and between-species in terms of Spearman correlation coefficient. Density plot of Spearman correlation between predicted and observed gene expression values across each species. (a) Distribution of the mean Spearman correlations for orthogroup expression predictions within each species. (b) Distribution of mean Spearman correlations for maize orthogroup expression predictions from *O. sativa, S*.*bicolor*, and the combination of *O. sativa* and *S*.*bicolor*.

We aim to maximize the covariance of *T* and *U* so that we can perform a regression such that

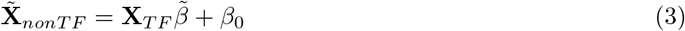

To find the optimal number of components cross-validation was performed, resulting in choosing 3 components.

### Graph autoencoder with random forest model

To create a low-dimensional representation of the expression and network data, we use a standard non-variational graph autoencoder, as seen in [29]. In this method, we consider input expression **X**_*tot*_ as a matrix with *N* genes and *Q* experiments and a graph *G*, as the Arabidopsis orthogroup iGRN. The encoder block of the autoencoder maps the regulatory network and gene expression features into a latent embedding (Fig 1). Then, in order to determine the validity of the embedding, the model decodes this embedding back into the input space and the cross-entropy loss of the reconstruction is measured. The predicted expression values are obtained using this latent representation **Z** such that

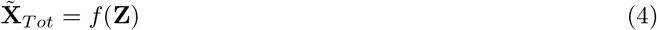

where 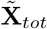 is the predicted matrix of the input expression values and f denotes a random forest with a max depth scaled by ⌊*Q*⌋ with ⌊⌋ denoting the floor operation.

To establish performance within this framework, we consider the use of two major encoding schemas, induction-based and convolution-based methods. For the induction-based method, we use the GraphSAGE layer available in PyTorch Geometric [41]. The GraphSAGE algorithm works by propagating hidden representations of the nodes (genes) to their neighbors [31]. Here, by stochastic descent, the combination of weight matrices and the mean aggregation function is tuned so that genes with similar behavior have similar representations within the resulting embedding [31]. Thus, a node, *i*, representation, *h*, for a neighborhood size of *k* is given by,

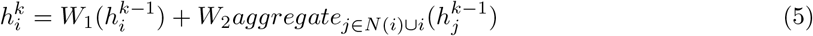

where the *aggregate* function is given by mean pooling and *W*_1_ and *W*_2_ are trainable weight matrices.

The convolutional framework takes into account the high order of structure in neighborhoods around nodes through the use of convolutional layers [43]. Hidden representations of those high-order structures are then merged together to form each node’s hidden state [43]. A single layer can be seen as

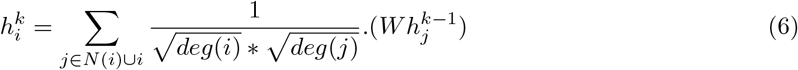

where *W* is a trainable weight matrix, *deg*(*i*) and *deg*(*j*) are the number of neighbors of node *i* and *j* respectively [43]. For the shallow autoencoder, we choose to use a single convolutional layer with hidden dimension of size 32. For the deep autoencoder, we choose three convolutional layers with a hidden dimension size of 32.

### Model evaluation

For each model task, we will use the Spearman rank correlation to compare the expression level of the observed and predicted orthogroup. Spearman’s is given by:

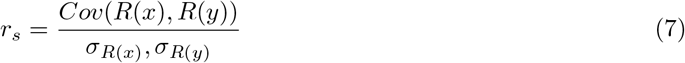

where *Cov*(*R*(*x*), *R*(*y*)) represents the covariance of the x and y ranks and *σ*_*R*(*x*)_,*σ*_*R*(*y*)_ represents the standard deviation of x *R*(*x*) and *R*(*y*), respectively.

### Network statistics

To understand the predictions with respect to structural embedding, we looked at three key network statistics: Degree centrality [30], Kleinberg Authority [30], and Harmonic centrality [44]. Degree centrality directly measures the total amount of edges connected to an individual node [30]. That is, for node *i* in the graph *G*, degree centrality is given by the degree of *i*:

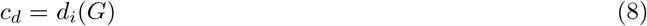

The harmonic centrality measure extends the degree of centrality by taking into account the distance between node i and all other nodes. To mitigate the effects of extreme high and low distances, the harmonic centrality calculation aggregates the inverse distances between nodes [30]. That is, for two nodes *i, j*:

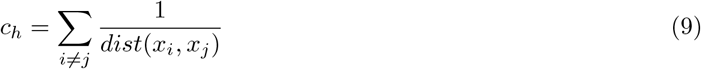

Finally, to examine the hub-like nature of each node, we calculated the Kleinberg Authority metric.

Here, we take the first eigenvector of **A**^*T*^ **A**, where **A** is the adjacency matrix of graph *G*. The eigenvalues of this vector gives a measure of how much information a node has (authority) and how well-connected the node is to other informative nodes (hubness) [44].

## Results

### Prediction data and model parameters

To predict the abundances of gene expression between experiments, we collected RNA-seq from *Sorghum bicolor, Oryza sativa*, and *Zea mays* and an *Arabidopsis thaliana* gene regulatory network. For the gene regulatory network, we took advantage of the *Arabidopsis thaliana* integrated gene regulatory network (iGRN) [35]. This network consists of 1,491 transcription factors and 31,393 target genes (about 27,000 protein-coding and a mixture of long noncoding RNAs and replicated transposable elements), resulting in a total of 1.9 million interactions weighted by the strength of a biologically supported interaction [35]. In order to reduce the number of parameters model estimates, we thresholded the network edges to only include an edge if its weight is at least 0.9. This gave a resulting network of about 250,000 interactions. For gene expression abundances, we used log2-transformed gene expression levels (in transcripts per million) that were collected in *Sorghum bicolor, Oryza sativa*, and *Zea mays*. We collapsed genes into orthogroups to allow for cross species comparisons. Table 1 describes a summary of the number of orthogroups, experiments, and transcription factors for each species.

**Table 1.** Summary of Input Data for Gene Expression Prediction Models.

| Species | Number of Transcription Factors | Number of Orthogroups | Number of Experiments |
| --- | --- | --- | --- |
| *Zea Mays* | 3336 | 18449 | 454 |
| *Sorghum bicolor* | 2665 | 16961 | 99 |
| *Oryza sativa* | 2409 | 20538 | 147 |

To set up a baseline for comparison with graph networks, we predicted non-transcription factor expression abundance from transcription factor expression using a partial least squares regression (Fig1). For the graph neural networks, we predicted expression under a graph autoencoder framework (Fig 1) with three different strategies for encoding the graph into a latent space. The first two strategies, deep and shallow convolutions, utilize graph convolutional layers, and the third strategy, GraphSAGE, uses an inductive convolutional layer that finds a mean representation of a node based on the information of nodes in its neighborhood. The representation of these models is then passed through a random forest model to predict gene expression profiles under different conditions. The models were tested in four scenarios: 1) prediction within species of unseen nonTF expression profiles when all genes have been seen by the model, 2) prediction in maize from a single species when all genes have been seen by the model, 3) prediction of unseen expression profile when all genes have not been seen by the model, and 4) prediction of maize expression profiles from all species.

### Partial least squares baseline performance

Before using the gene regulatory network model, we performed a partial least squares regression with three components. The partial least squares model does not use any information from the gene regulatory network’s structure. For each species, we predicted the expression profile of 1434 genes across 16 experiments and compared those to the observed abundances. To assess the performance of the model we decided to use Spearman’s rank correlation. *S. bicolor* performed the best at a correlation of mean 0.80, but *Z. mays* (*ρ* = 0.77) and *O. sativa* (*ρ* = 0.76) correlated well. Figure 3a gives the distribution of the mean correlations experiments with respect to each species.

**Fig 3.**
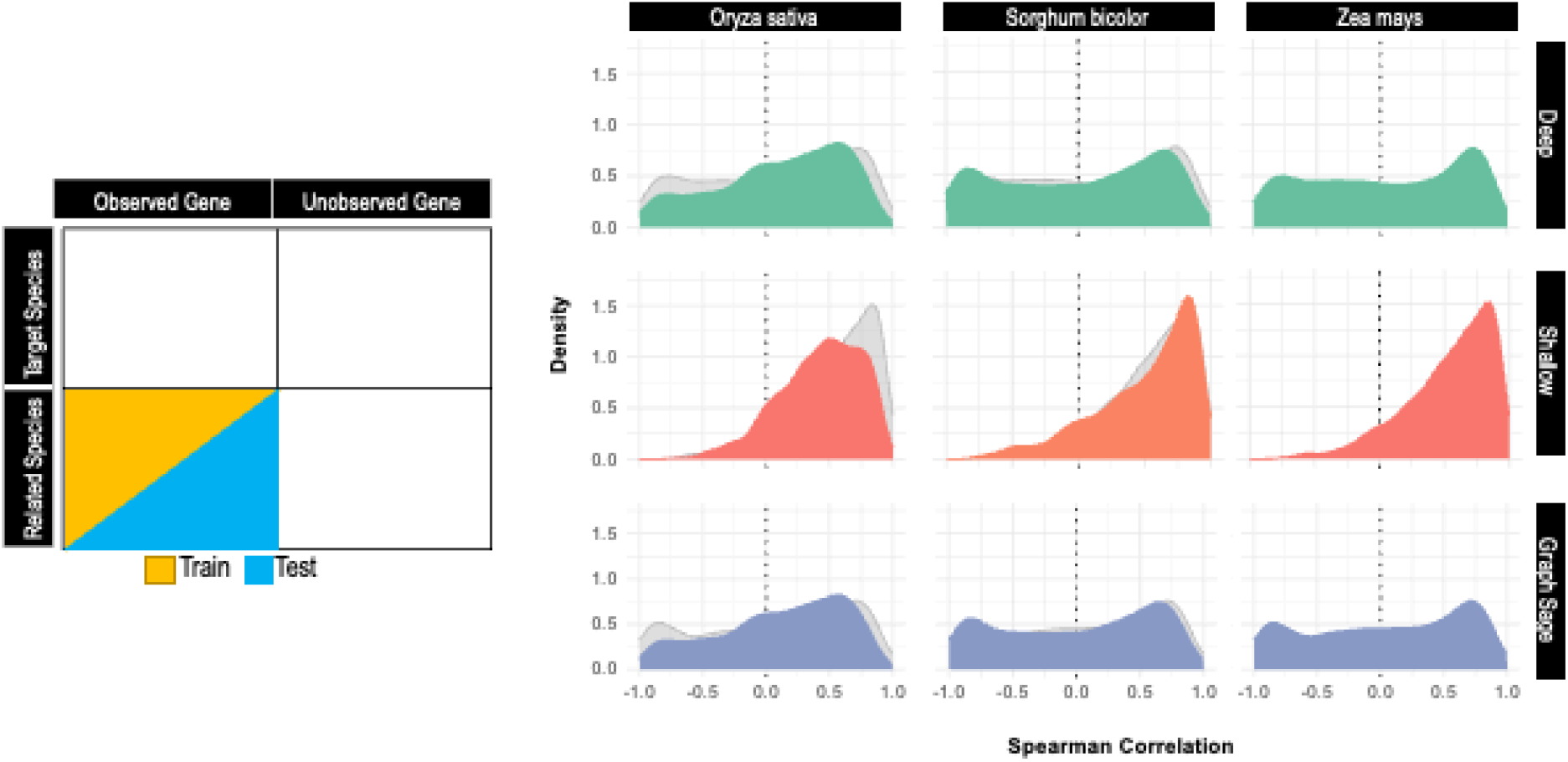
Prediction performance of within-species autoencoder models in terms of the Spearman correlation coefficient. Density plot of Spearman correlation between predicted and observed gene expression values across each species. Three strategies for mapping to latent spaces, Shallow Graph Convolutions, Deep Convolutions, and Graph Sage were utilized across *Oryza sativa, Sorghum bicolor*, and *Zea mays*.

For the next baseline, we wanted to see how well each species does to naively predict maize ortholog expression. To do this, we separated all of the maize ortholog expressions from all of the other species. Next, we used each species’ ortholog profiles to predict the corresponding ortholog’s profile. While overall lower than maize just predicting within itself, we found that the average Spearman correlations were still comparable to within species predictions. When we used *S. bicolor* to predict maize, we get correlation of 0.77. Similarly, when we used *O. sativa* to predict maize expressions we got correlation of 0.74, we used wanted to look at how well PLS does when we are using *O. sativa* and *S. bicolor* orthologs to predict maize expression abundance. For the cases when we used *O. sativa* and *S. bicolor*, we find that the mean Spearman correlations are 0.74 and 0.78, respectively. These results show that regardless of species combination, partial least square models tend to estimate gene expression similarly between and within species.

### Graph autoencoder model performance not comparable for within-species predictions

To test the hypothesis that graph structure would improve baseline predictions, we implemented a graph autoencoder (GAE) using three different encoder architectures. We trained each model within *Oryza sativa, Sorghum bicolor*, and *Zea mays* to predict expression profiles within each species. For each species. the training sets were 75% of each species’ orthogroups. To ensure, no data leakage, we masked the 25% test set during the training process. We ran each model for a total of 200 epochs with a learning rate of 0.001. The first method, Deep GAE, performed poorly in all species with mean Spearman correlations of 0.116, 0.179, 0.117 for *Oryza sativa, Sorghum bicolor*,and *Zea mays*, respectively. The inductive method, GraphSage, also did not perform well within species with mean correlations of 0.054, 0.052, and -0.015. The best performing method, the shallow graph autoencoder, gave mean predictions of 0.46 in *Oryza sativa*, 0.56 in *Sorghum bicolor* and 0.61 in *Zea mays*. Figure 3 shows the distributions of the Spearman correlations of the predictions.

### Shallow graph autoencoders outperform complex strategies in cross-species expression predictions

We next predicted in unseen maize gene expressions using the best performing strategy-the shallow graph autoencoder. We used the shallow graph convolutional autoencoder in three models covering *Z. mays + O. sativa, Z. mays + S. bicolor*, and *Z. mays + O. sativa + S. bicolor*. Again, we used the transcription factor expressions as input to the model to predict the unobserved maize gene expression profiles. Each model predicts 1435 unobserved maize genes across 217 experiments.In all three contexts, the shallow graph allowed the predictions of these unseen genes in maize with *ρ* of 0.42 to 0.47 (Fig 4).

**Fig 4.**
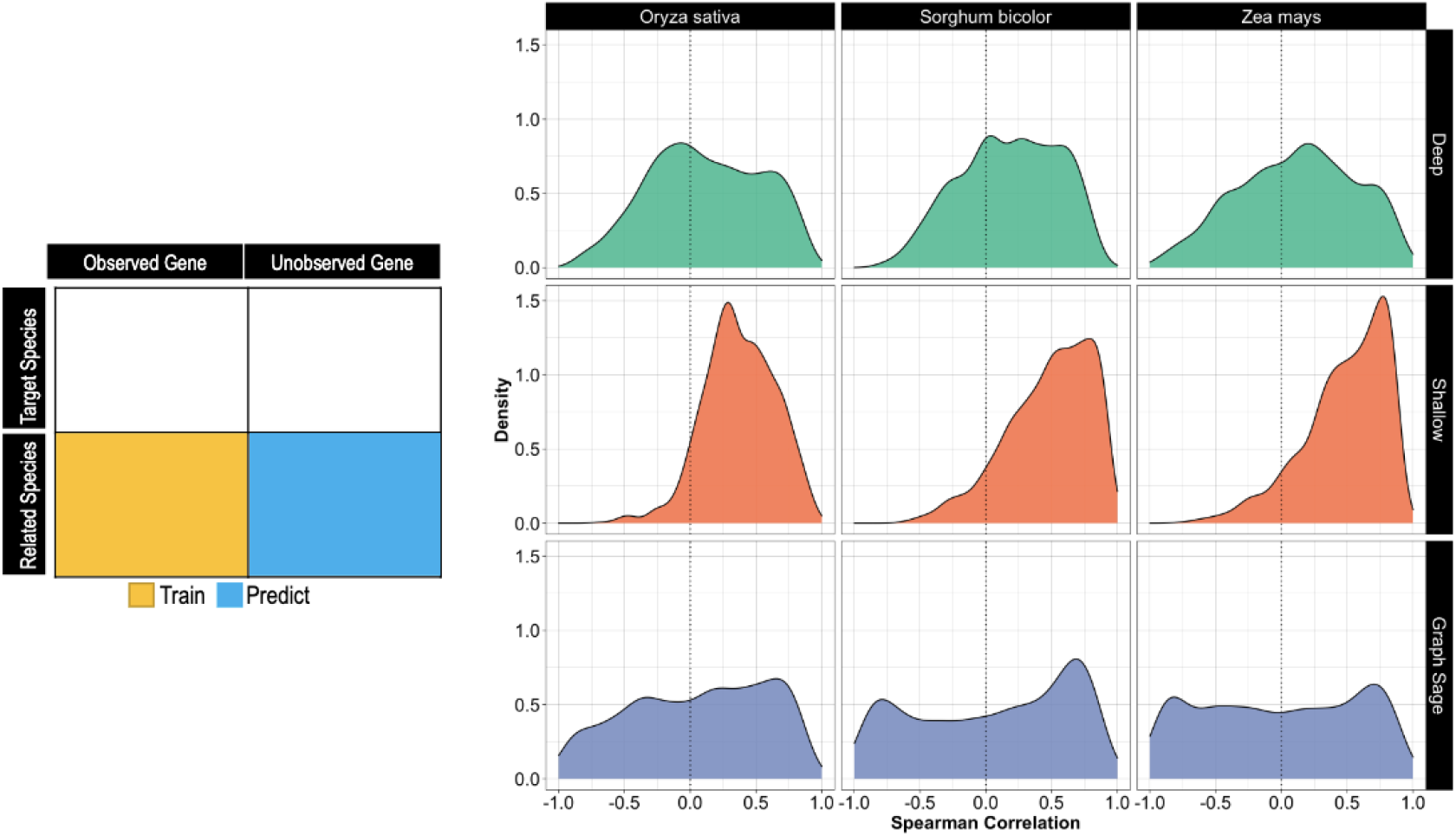
Distribution of the predictive performance of the shallow graph convolutional autoencoder for the abundance of maize expression. Density plot of predictive performance Spearman Correlation of graph convolutional autoencoder. Predictive performance (x-axis) is given as the Spearman Correlation between observed and predicted orthogroup expressions. Plots are colored according to the species at hand. Dashed, vertical lines represent the mean of each distribution.

### Graph autoencoders allow for predictions in never-before-seen samples

To test the utility of using a graph autoencoder for gene expression predictions, we predicted gene expression in genes which were previously unobserved in the training. To do this, we first made sure that any genes the model could possibly predict would be included are accounted for in the iGRN. Next, we used the three autoencoder strategies to predict unobserved genes in masked experiments within each species. We trained each model for 2000 epochs with a learning rate of 0.001. Just as with the within species model, we note that the GraphSage and Deep autoencoder strategies performed the worst. The deep autoencoder had within-species prediction performances for *O. sativa, S. bicolor*, and *Z*.*mays* of -0.10, 0.31, and 0.11, respectively. GraphSage had correlations of 0.01, 0.02, and 0.002 for *O. sativa, S. bicolor*, and *Z*.*mays*, respectively. The shallow autoencoder, however, performed surprisingly well compared to the observed gene, within-species baselines with correlations of 0.47 - 0.53 (Fig 5).

**Fig 5.**
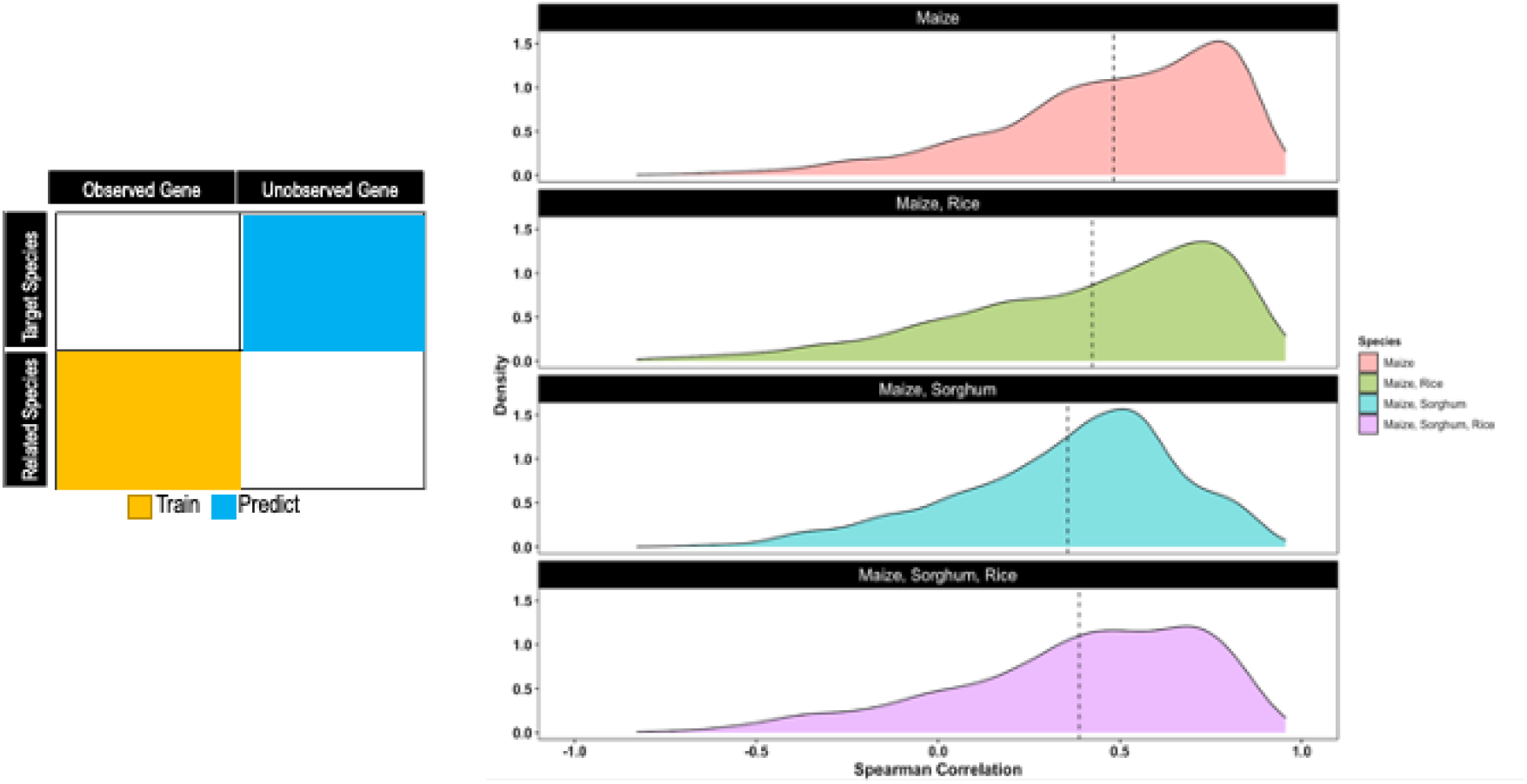
Distribution of predictive performance of shallow graph convolutional autoencoder for maize expression abundance. Density plot of predictive performance Spearman Correlation of graph convolutional autoencoder. Predictive performance (x-axis) is given as the Spearman Correlation between observed and predicted orthogroup expressions. Plots are colored according to the species at hand. Dashed, vertical lines represent the mean of each distribution.

**Fig. 6.**
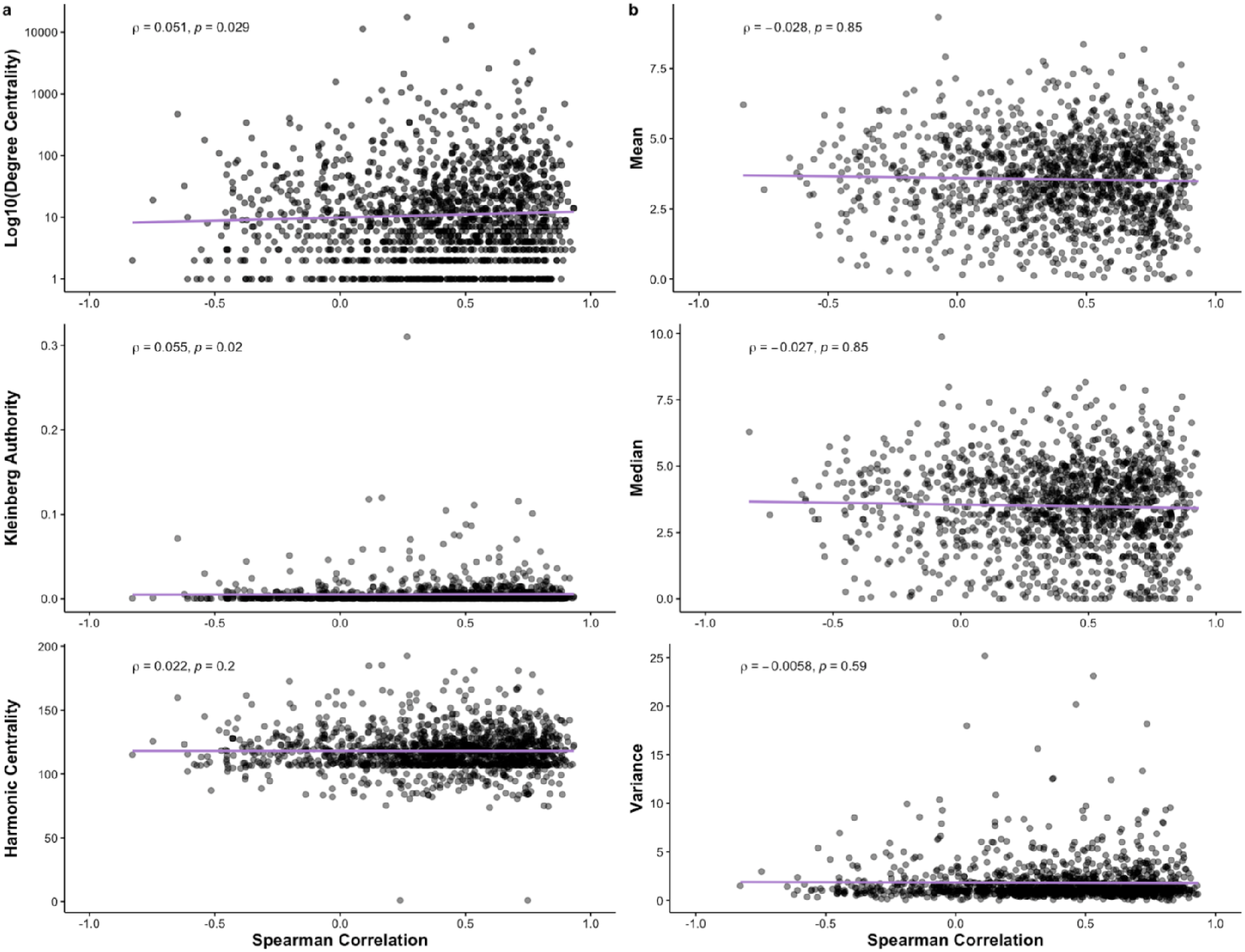
Relationships between shallow graph convolutional autoencoder performances and key descriptive statistics. (a) Relationship between three common network measures, Degree Centrality, Kleinberg Authority, and Harmonic Centrality, with the Spearman correlations for predicting maize expression abundance with a shallow graph autoencoder. (b) Relationship between descriptive statistics about the expression data, mean, median, and variance, with the Spearman correlations for predicting maize expression abundance with a shallow graph autoencoder.

### Predictive performance is not strongly driven by network structure or variation in expression

Our final hypothesis was that structure of the Arabidopsis iGRN and the intensity of gene expression drives model prediction accuracy. In order to test the hypothesis, we plotted the maize cross-species prediction accuracy against network descriptive statistics and expression statistics. (Figure 4a) and expression statistics (Figure 5). In the case of the network statistics, there was little to no negative correlation between the measure of Log10-centrality (*ρ* = 0.051) and Kleinberg Authority (*ρ* = 0.055) and harmonic centrality (*ρ* = 0.022). These findings indicate that the influence of network structure on the model is minimal. For the expression variation statistics, we use Spearman correlation to compare overall gene expression mean, median, and variance with predictive performance. Just as with network statistics, we find there is a very slightly negative relationship between mean (*ρ* = −0.028), median (*ρ* = −0.027), and variance (*ρ* = −0.0058), but we note that these values are not significant.

## Discussion

Gene expression inference across species, experiments, and time require effective technological frameworks, carefully curated data, and powerful mathematical frameworks. In traditional gene expression inference frameworks, such as the LINCS linear regression-based platform, there is little focus on the non-linearity of gene expression. We present a way to utilize the gene regulatory network’s nonlinear connections in a prediction framework that allows for the prediction of gene expression values for unobserved genes in unobserved experiments and is comparable to within-species predictions. Our results suggest that gene expression prediction accounting for discrete biological network structure can be used to achieve moderate-accuracy target gene expression values.

We note that there is a large imbalance between the accuracy of predictions in the within-species models. In *Sorghum bicolor*, this discrepancy is likely to be from a few genes which are highly variable.

In *Zea mays* and in *Oryza sativa*, our hypothesis is that this discrepancy is likely due to the different types of tissue classes included in the dataset. There is evidence that the transcription factor background can be attributed to the determination of biological roles such as tissue specificity [45], gene function [17], and cellular behavior [46]. In addition to the overall differences per species, there is a wide range of differences between each of the model types.

One of the main results of the study is that the shallow convolution operator consistently outperforming the more complex models. We hypothesize this is due to a combination of the network structure and the ability for simpler model to generalize on new data. For the network case, we decided to take the most representative edges in the Arabidopsis iGRN. Deeper GNNs and GraphSAGE models inform their embedding projections by taking in information from surrounding nodes. By paring down the network at hand, these models put heavy emphasis on nodes that are already known to have a large bias toward genes that are known and expressed. This means that the 2nd and 3rd order relationships that may be in the larger, less thresholded network are largely unseen. This can lead to under-accounting for certain relationships and over-accounting for relationships that actually involve more connections. The relationships at question, however, must generalize across species and expression pattern. As seen in genomic prediction and language modeling, the less complex model often performs better than the more complex models [47].

One of the key problems in this analysis is that we only use one data modality, RNA-seq. The landscape of gene expression is rather complex and requires not only understanding the non-linearity of regulator-target relationships, but also understanding the interactions between regulatory elements that cannot be identified exclusively in assays such as RNA-seq.This type of modeling has been done with models such as sequence-specific models which use wide ranges of regulatory information to guide the model’s training and testing. In other models, such as GC-merge [48], cis-regulatory information from Chip-Seq integrates expression measurements with a deep graph convolutional framework much like the one we utilized. In the future, prediction across species using more types of readily-accessible data could provide a more biologically informed way to power these sophisticated deep-learning models.

## Supporting information

**S1 Fig.**
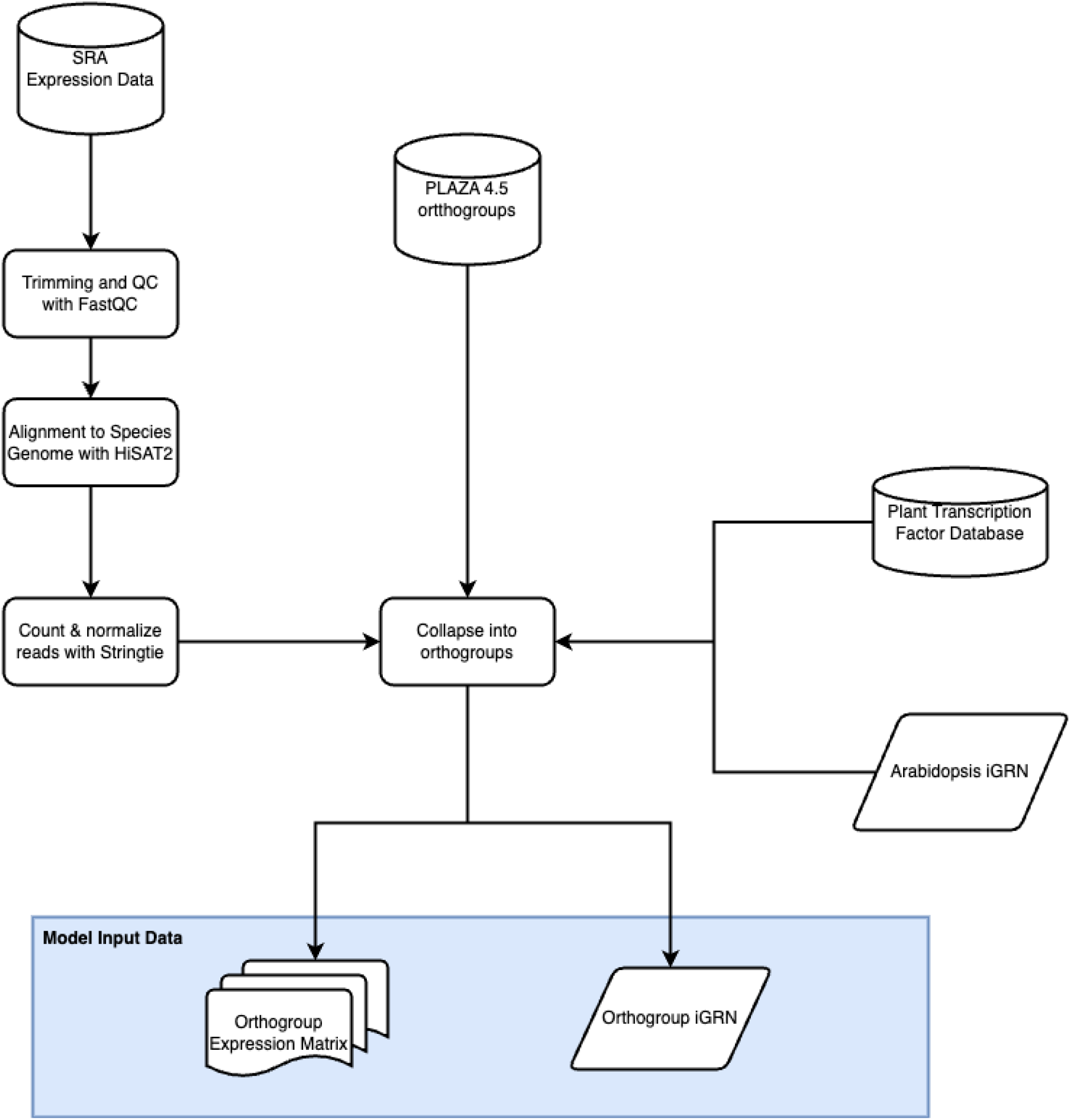
Flowchart for pre-processing of model input data.

## Acknowledgments

We thank Travis Wrightsman and Michelle Stitzer for their suggestions in the preparation of this manuscript.

## Notes

### Competing Interest Statement

The authors have declared no competing interest.

